# HDAC1/2 and HDAC3 play distinct roles in controlling adult Meibomian gland homeostasis

**DOI:** 10.1101/2024.02.06.579230

**Authors:** Xuming Zhu, Mingang Xu, Sarah E. Millar

**Author notes:** Author for correspondence: Sarah E Millar.

## Abstract

**Purpose:** To investigate the roles of HDAC1/2 and HDAC3 in adult Meibomian gland (MG) homeostasis.

**Methods:** HDAC1/2 or HDAC3 were inducibly deleted in MG epithelial cells of adult mice. The morphology of MG was examined. Proliferation, apoptosis, and expression of MG acinus and duct marker genes, meibocyte differentiation genes, and HDAC target genes, were analyzed via immunofluorescence, TUNEL assay, and RNA in situ hybridization.

**Results:** Co-deletion of HDAC1/2 in MG epithelium caused gradual loss of acini and formation of cyst-like structures in the central duct. These phenotypes required homozygous deletion of both HDAC1 and HDAC2, indicating that they function redundantly in the adult MG. Short-term deletion of HDAC1/2 in MG epithelium had little effect on meibocyte maturation but caused decreased proliferation of acinar basal cells, excessive DNA damage, ectopic apoptosis, and increased p53 acetylation and *p16* expression in the MG. By contrast, HDAC3 deletion in MG epithelium caused dilation of central duct, atrophy of acini, defective meibocyte maturation, increased acinar basal cell proliferation, and ectopic apoptosis and DNA damage. Levels of p53 acetylation and p21 expression were elevated in HDAC3-deficient MGs, while the expression of the differentiation regulator PPARγ and the differentiation markers PLIN2 and FASN was downregulated.

**Conclusions:** HDAC1 and HDAC2 function redundantly in adult Meibomian gland epithelial progenitor cells and are essential for their proliferation and survival, but not for acinar differentiation, while HDAC3 is required to limit acinar progenitor cell proliferation and permit differentiation. HDAC1/2 and HDAC3 have partially overlapping roles in maintaining survival of MG cells.

## Introduction

Dry Eye Disease (DED) is prevalent, affecting almost 7% of the US adult population. Untreated DED can lead to inflammation of the eye, corneal abrasions and ulcers, and vision loss [1, 2]. The most common type of DED, known as evaporative DED (EDED), is associated with decreased production of meibum, an oily film that covers the eye surface and prevents tear film evaporation [3, 4]. Decreased meibum production results from dysfunction of the Meibomian glands (MGs) [1]; these are specialized sebaceous glands located in the tarsal plate of the upper and lower eyelids whose major function is to secrete meibum [3, 4]. MG shrinkage or dropout, accompanied by reduced quantity and quality of meibum, is increased in aging; in line with this, EDED incidence is highest in aging populations [5, 6]. Available treatments, including lipid-containing eye drops, warm compresses to improve meibum flow, and anti-inflammatory drugs, provide limited and temporary efficacy for EDED, and the disease currently has no cure [7]. Improved understanding of the cellular and molecular processes underlying MG homeostasis will be crucial in developing more effective therapeutics for this common condition that negatively affects quality of life.

Each MG consists of a central duct and multiple connecting acini, in which basal stem cells constantly proliferate toward the center of acinus to produce meibocytes that undergo sequential differentiation to accumulate lipid content for meibum synthesis [8, 9]. Failed MG stem cell activity is proposed as an underlying cause of MG dropout [10, 11], but the mechanisms that govern the proliferation, survival, and differentiation of MG stem cells are poorly understood.

Chromatin regulators play key roles in governing stem cell proliferation, differentiation, and maintenance in epithelial tissues, and are druggable targets with therapeutic potential. Among them, histone deacetylases (HDACs) remove histone acetyl groups, resulting in compaction of chromatin and transcriptional repression [12, 13]. HDACs can also deacetylate key transcription factors, such as p53 and c-MYC, regulating their stability and activity [14, 15].

The Class I HDACs, HDAC1, HDAC2, and HDAC3, have strong deacetylase activity towards histones and are associated with at least four major repressive complexes, indicating that they have isoform and context-dependent effects. HDAC1 and HDAC2 are closely related, function redundantly, or semi-redundantly, in some contexts [16, 17], and associate with the nucleosome remodeling and deacetylase complex (NuRD) [18, 19], the transcriptional regulatory protein Sin3A [20], corepressor of REST (CoREST) complex [21], and the mitotic deacetylase complex (MiDAC) [22]. By contrast, HDAC3 uniquely associates with the SMRT/N-CoR corepressor complex [23], suggesting that it regulates a set of target genes distinct from that controlled by HDAC1/2. In line with this, HDAC1/2 are required for proliferation and survival of epidermal basal cells [24, 25], while HDAC3 plays essential roles in preventing premature epidermal differentiation [26].

HDAC inhibitors have been developed and approved for the treatment of cancers such as cutaneous and peripheral T-cell lymphoma, as well as multiple myeloma [27, 28]. HDAC inhibitors can also have anti-inflammatory effects [29, 30]. As DED is an inflammatory ocular disease [31, 32], HDAC inhibitors have been suggested as an alternative therapeutic approach for DED. This notion gains support from a study showing that controlled release of an HDAC inhibitor into the lacrimal gland can effectively mitigate inflammation in DED-afflicted mice [33]. However, currently available HDAC inhibitors generally act broadly on multiple HDAC family members and can have severe side effects [34, 35]. Designing more specific therapeutics will require improved knowledge of the functions of individual HDACs in MG homeostasis. While constitutive deletion of *Hdac1* and *Hdac2* in developing MG of *K14-Cre Hdac1^fl/fl^ Hdac2^fl/+^* mice causes MG hyperplasia [36], the functions of *Hdac1* and *Hdac2* in adult MG epithelium have not been examined, and any roles played by *Hdac3* in this tissue are unknown. To investigate the overlapping and unique functions of HDAC1/2 and HDAC3 in MG homeostasis we examined the phenotypic and molecular consequences of inducible deletion of these genes in adult MG epithelium.

## Materials and methods

### Mice

The following mouse lines were used: *Hdac1^fl^* and *Hdac2^fl^* [37]; *Hdac3^fl^*[38]; *Krt5-rtTA* [39]; and *tetO-Cre* [40]. Mice were allocated to experimental or control groups according to their genotypes, with littermate mice lacking *Krt5-rtTA* or *tetO-Cre* used as controls in each experiment. Mice were included in the analysis based on their genotypes. Investigators were aware of genotype during allocation and animal handling as this information was required for appropriate allocation and handling. Up to five mice were maintained per cage in a specific pathogen-free barrier facility on standard rodent laboratory chow (Purina, 5001). To induce DNA recombination, mice were fed with doxycycline (Sigma-Aldrich, 200µg/mL in 5% sucrose) for the indicated time. Mice were maintained on a mixed strain background and all animal experiments were performed under approved animal protocols according to institutional guidelines established by the Icahn School of Medicine at Mount Sinai IACUC committee. At least three mice of each genotype and experimental condition were assayed for all the experiments in this study.

### Histology, immunofluorescence (IF), and TUNEL assays

Tissues were fixed in 4% paraformaldehyde (PFA)/PBS solution (Affymetrix/USB) overnight at 4°C, embedded in paraffin, and sectioned at 5 µm. Sections were de-paraffinized and rehydrated through xylene substitute (Sigma-Aldrich) and graded ethanol solutions. Sections underwent antigen retrieval in the unmask solution (Vector Labs). For IF, sections were incubated with primary antibodies overnight at 4°C, followed by incubation with fluorescently labeled secondary antibodies (Invitrogen and Vector Labs) and mounting medium (Invitrogen). For TUNEL assays, rehydrated sections were incubated with TUNEL labeling mix (Roche) at 37°C for 1 hour, washed with PBS+0.1% Tween 20 (PBST), and mounted with DAPI-containing mounting medium (Invitrogen).

### Antibodies

The following primary antibodies were used: rabbit anti-HDAC1 (Thermo Fisher Scientific, 49-1025, 1:1000); rabbit anti-HDAC2 (Thermo Fisher Scientific, 51-5100, 1:1000); rabbit anti-HDAC3 (Santa Cruz, sc-11417, 1:100); rabbit anti-PLIN2 (Fitzgerald, 20R-AP002, 1:500); rat anti-CD45 (BioLegend, 50-162-774, 1:100); rabbit anti-p21 (Proteintech, 28248-1-AP, 1:200); rabbit anti-KRT6A (BioLegend, 905702, 1:200); mouse anti-KRT14 (Invitrogen, #MA5-11599, 1:500); rat anti-Ki-67 (eBioscience, 14-5698-82, 1:200); rabbit anti-KRT5 (Covance, PRB-160P, 1:1000); rabbit anti-acetyl-p53 (Abcam, ab61241, 1:200); rabbit anti-KRT10 (Covance, PRB-159LP, 1:1,000); rabbit anti-PPARγ (Cell Signaling, 2435T, 1:200); rabbit anti-FASN (Cell Signaling, 3180T, 1:200); rabbit anti-KRT17 (Abcam, ab109725, 1:200); rabbit anti-pH2A.X (Cell Signaling, #9718, 1:200).

The following secondary antibodies were used: donkey anti-rabbit IgG, Alexa Fluor 555 (Invitrogen, A-31572, 1:800); donkey anti-mouse IgG, Alexa Fluor 488 (Invitrogen, A-21202, 1:800); goat anti-rabbit IgG, Alexa Fluor 488 (Invitrogen, A-11008, 1:800); goat anti-mouse IgG, Alexa Fluor 555 (Invitrogen, A-28180, 1:800); goat anti-rat IgG, Alexa Fluor 488 (Invitrogen, A-11006, 1:800); goat anti-guinea Pig, Biotinylated (Vector Labs, BA-7000, 1:500); Streptavidin Fluorescein (Vector Labs, SA-5001, 1:500); Streptavidin Texas Red (Vector Labs, SA-5006, 1:500);

### RNAscope

RNAscope was performed on paraffin sections following the protocol provided by Advanced Cell Diagnostics (ACD) using RNAscope Multiplex Fluorescent Reagent Kit v2 (ACD #323100) and probes for *Mdm2* (ACD #447641), *p16* (ACD #411011), *Ccnd1* (ACD #442671), *Ptch1* (ACD #402811) and *Gli1* (ACD #311001-C2). The sections were observed and photographed using a Leica Microsystems DM5500B fluorescent microscope.

### Statistical analysis

Statistical analysis and graphical representation were performed using Microsoft Excel 2023. Unpaired two-tailed Student’s *t*-test was used to calculate statistical significance between two groups of datasets. *P* < 0.05 was considered significant.

## Results

### HDAC1/2 function redundantly to maintain adult MG homeostasis

To begin to explore the roles of HDAC1 and HDAC2 in adult MG homeostasis, we first examined their expression in adult MG. IF staining revealed that HDAC1 exhibits broad expression in MG acini and central ducts, but is undetectable in mature meibocytes (Figure 1A, white arrow). HDAC2 is expressed more broadly than HDAC1, including in mature meibocytes (Figure 1B, yellow arrow). To determine the functions of *Hdac1* and *Hdac2* in adult MG, we generated *K5-rtTA tetO-Cre Hdac1^fl/fl^ Hdac2^fl/+^*(*Hdac1^cKO^ Hdac2^cHet^*) mice in which doxycycline administration triggers homozygous *Hdac1* deletion and heterozygous deletion of *Hdac2* in adult MG epithelial cells. Similarly, we generated *K5-rtTA tetO-Cre Hdac1^fl/+^ Hdac2^fl/fl^* (*Hdac1^cHet^ Hdac2^cKO^*) mice in which homozygous *Hdac2* deletion and heterozygous deletion of *Hdac1* in MG epithelium is induced by doxycycline treatment (Fig. S1A). MG histology in both mutants was indistinguishable from that of controls after two weeks of doxycycline treatment (Figure S1B-D) despite the absence of HDAC1 or HDAC2 in MG epithelium of *Hdac1^cKO^ Hdac2^cHet^* and *Hdac1^cHet^ Hdac2^cKO^* mice, respectively (Fig. S1E-J). MG duct structure and meibocyte differentiation were similar in mutant and control mice, as indicated by normal expression of the MG duct marker KRT6A and the meibocyte differentiation marker PLIN2 [9] (Fig. S1K-M). Thus, while constitutive homozygous *Hdac1* deletion paired with heterozygous *Hdac2* loss in the developing MG is sufficient to cause MG hyperplasia [36], these manipulations do not appear to perturb the homeostasis of the established adult MG.

**Figure 1.**
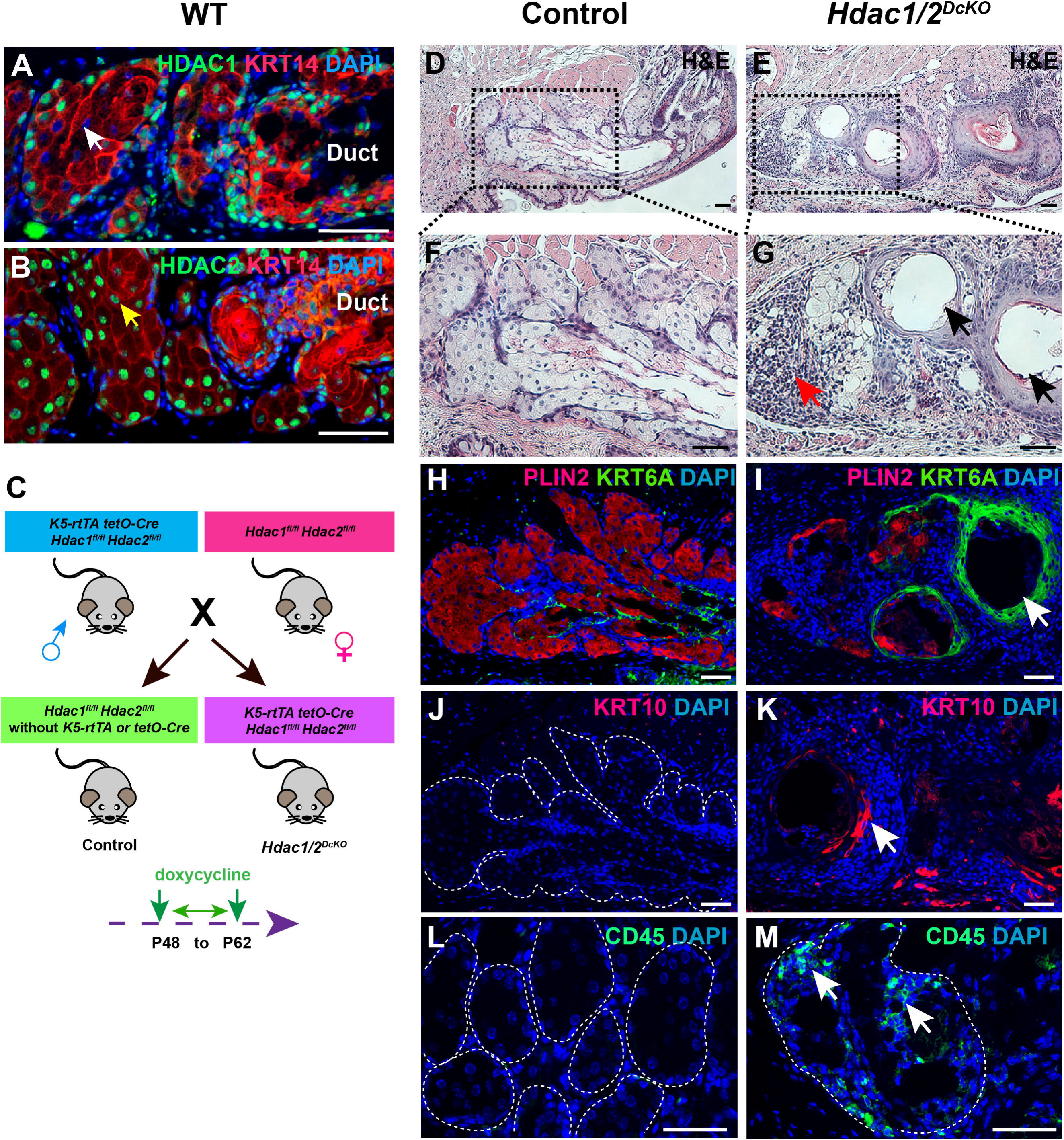
Co-deletion of HDAC1/2 in adult MG epithelium causes MG degeneration. (A) IF staining reveals broad expression of HDAC1 in KRT14+ MG epithelial cells, and its absence in mature meibocytes (white arrow) in wild-type (WT) mice. (B) IF staining shows ubiquitous expression of HDAC2 in WT MG epithelium, including in mature meibocytes (yellow arrow). (C) Schematic representation of the mouse breeding strategy and experimental design. (D, E) H&E staining shows normal structural organization of control MG (D) and degeneration of MGs in *Hdac1/2^DcKO^* mice (E). (F, G) Amplified views of the outlined regions in (D) and (E), respectively. Arrows in (G) highlight infiltrative cells in the acinus (red arrow) and cyst-like structures apparently originating from the MG duct (black arrows). (H) IF analysis of control MG shows KRT6A expression predominantly in MG duct and PLIN2 expression in meibocytes. (I) IF indicates fewer numbers of PLIN2+ meibocytes and the presence of KRT6A+ cyst-like structures (white arrow) in HDAC1/2-deficient MGs. (J, K) IF shows absence of KRT10 expression in control MGs (J) and ectopic KRT10 expression in cyst-like structures in *Hdac1/2^DcKO^* MG (K, white arrow). (L, M) CD45+ immune cells are not evident in control MG (L) but infiltrate the degenerating MG acini in *Hdac1/2^DcKO^* MG (M, white arrows). White dashed lines delineate MG acini. Scale bars: 50μm.

These data suggested that *Hdac1* and *Hdac2* may function redundantly in the adult MG. To test this, we generated *K5-rtTA tetO-Cre Hdac1^fl/fl^ Hdac2^fl/fl^*(*Hdac1/2^DcKO^*) mice and subjected them to doxycycline treatment in adult life to determine the effects of simultaneous homozygous deletion of both *Hdac1* and *Hdac2* in adult MG epithelium (Fig. 1C). Following two-weeks of doxycycline induction, histological examination of MGs of *Hdac1/2^DcKO^* mice revealed signs of degeneration (Fig. 1D-G), including partial meibocyte loss, immune cell infiltration in the acini (Fig. 1G, red arrow), and formation of cyst-like structures in the MG duct (Fig. 1G, black arrows). IF results confirmed partial loss of PLIN2+ meibocytes (Fig. 1H and 1I). The cyst-like structures in HDAC1/2-deleted MG expressed the MG duct marker KRT6A suggesting that they may derive from the MG duct (Fig. 1H and 1I). These structures also exhibited expression of the epidermal differentiation marker KRT10 (white arrow, Fig. 1K), which is not expressed in normal MG ducts (Fig. 1J), indicating a shift in the fate of duct cells towards epidermis upon HDAC1/2 deletion. HDAC1/2-deleted acini contained CD45+ immune cells (white arrows, Fig. 1M), which were absent in control acini (Fig. 1L). Collectively, these findings reveal that HDAC1 and HDAC2 play essential, redundant roles in the maintenance of adult MGs.

### HDAC1/2 are essential to maintain MG basal cell proliferation and survival

To unravel the mechanisms through which HDAC1/2 exert their influence on MG homeostasis, we limited the doxycycline induction period to 10 days, a time point at which the structural integrity of HDAC1/2-deficient MGs remained relatively intact (Fig. 2A and 2B). IF analyses confirmed efficient ablation of HDAC1/2 within the MG epithelium at this stage (Fig. 2C and 2D). Notably, expression of the duct marker KRT6A, as well as the meibocyte markers PLIN2 and FASN, along with the meibocyte differentiation regulator marker PPARγ, was unaltered in HDAC1/2-deleted MGs compared with controls (Fig. 2E-J), indicating that HDAC1/2 are not required for meibocyte differentiation. However, within HDAC1/2-deficient MGs, acinar basal cell proliferation, assayed by IF for Ki-67, was significantly reduced (Fig. 2K, 2L, and 2Q). TUNEL assays and pH2A.X IF results showed that in the absence of HDAC1/2, MGs displayed ectopic apoptosis (Fig. 2M and 2N) and DNA damage (Fig. 2O and 2P) in the ducts (Fig. 2N and 2P, white arrows) and acini (Fig. 2N and 2P, yellow arrows). These data revealed key roles for HDAC1/2 in promoting acinar basal cell proliferation and ensuring the survival of cells within both the MG duct and the acinus.

**Figure 2.**
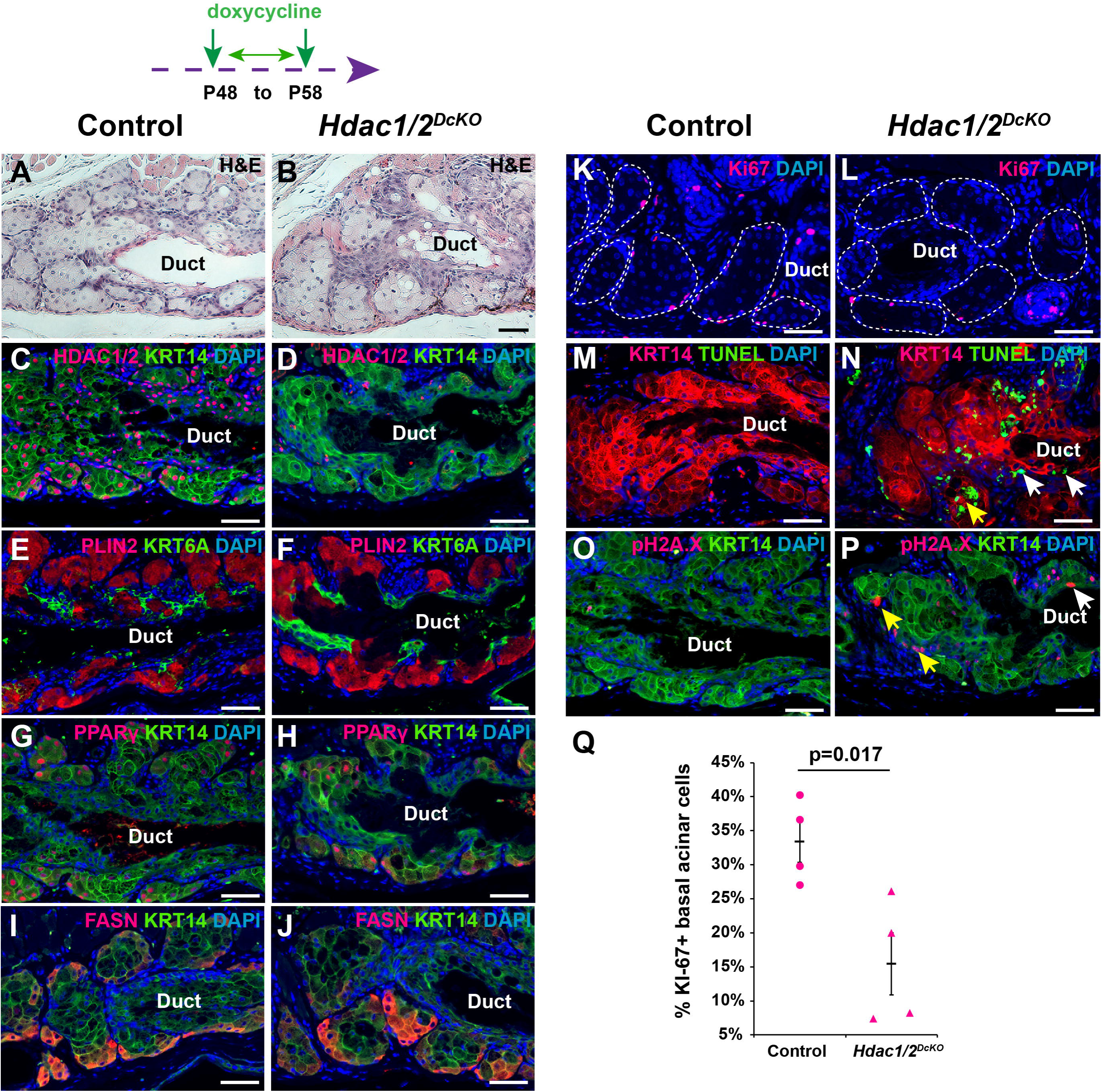
HDAC1/2-deficient MGs exhibit decreased acinar basal proliferation, ectopic apoptosis, and DNA damage. (A, B) H&E staining after 10 days of doxycycline treatment shows normal morphology of control (A) and aberrant morphology of *Hdac1/2^DcKO^* (B) MGs. (C, D) IF for HDAC1/2 shows their efficient deletion in KRT14+ MG epithelium of *Hdac1/2^DcKO^* mice subjected to a 10-day doxycycline treatment induction (D) compared with littermate control (C). (E, F) Expression of PLIN2 in meibocytes and KRT6A in the MG is unaffected by HDAC1/2 deletion (F) compared with littermate control (E). (G, H) Expression of PPARγ in KRT14+ MG epithelial cells is unchanged in Hdac1/2DcKO mice after 10 days of doxycycline treatment (H) compared with littermate control (G). (I, J) Expression of FASN is unchanged in *Hdac1/2^DcKO^*MG acini after 10 days of doxycycline treatment (J) compared with littermate control (I). (K, L) The number of Ki-67+ basal cells in the acinar basal layer of *Hdac1/2^DcKO^* mice after 10 days of doxycycline treatment (L) is reduced compared with littermate control (K). (M, N) TUNEL assays reveal minimal apoptosis in control MGs (M) and ectopic apoptosis in the acini (yellow arrow) and duct (white arrows) of HDAC1/2-deficient MGs (N). (O, P) IF for pH2A.X reveals absence of detectable DNA damage in basal cells of control MGs (O) and pronounced DNA damage in the basal layer of acini (yellow arrows) and duct (white arrow) in HDAC1/2-deficient MGs (P). (Q) Quantification of Ki-67 positive cells in the acinar basal layer of control and *Hdac1/2^DcKO^*MGs; n=4 controls and n=4 mutants. Error bars indicate SEM. White dashed lines in (K, L) delineate MG acini. Scale bars: 50μm.

### HDAC1/2 are required for p53 deacetylation and suppression of *p16* expression in adult MG

Previous studies showed that HDAC1/2 maintain basal cell proliferation and survival in the epidermis by limiting p53 acetylation and suppressing *p16* expression [24, 25]. To determine whether similar mechanisms operate in the adult MG, we assayed for acetylated p53 (p53Ac) in HDAC1/2-deficient MGs and discovered that this was significantly elevated (Fig. 3A and 3B). In line with this, IF and RNA in situ hybridization assays showed that expression of the p53 targets p21 and *Mdm2* was upregulated in HDAC1/2-depleted MGs (Fig. 3C-F, yellow arrows). RNAscope assays revealed elevated *p16* expression in HDAC1/2 mutant MGs compared with controls (Fig. 3G and 3H, yellow arrows). Together, these findings suggested that HDAC1/2 regulate the proliferation and survival of MG epithelial cells by repressing p53 acetylation and inhibiting *p16* expression, similar to their functions in the epidermis.

**Figure 3.**
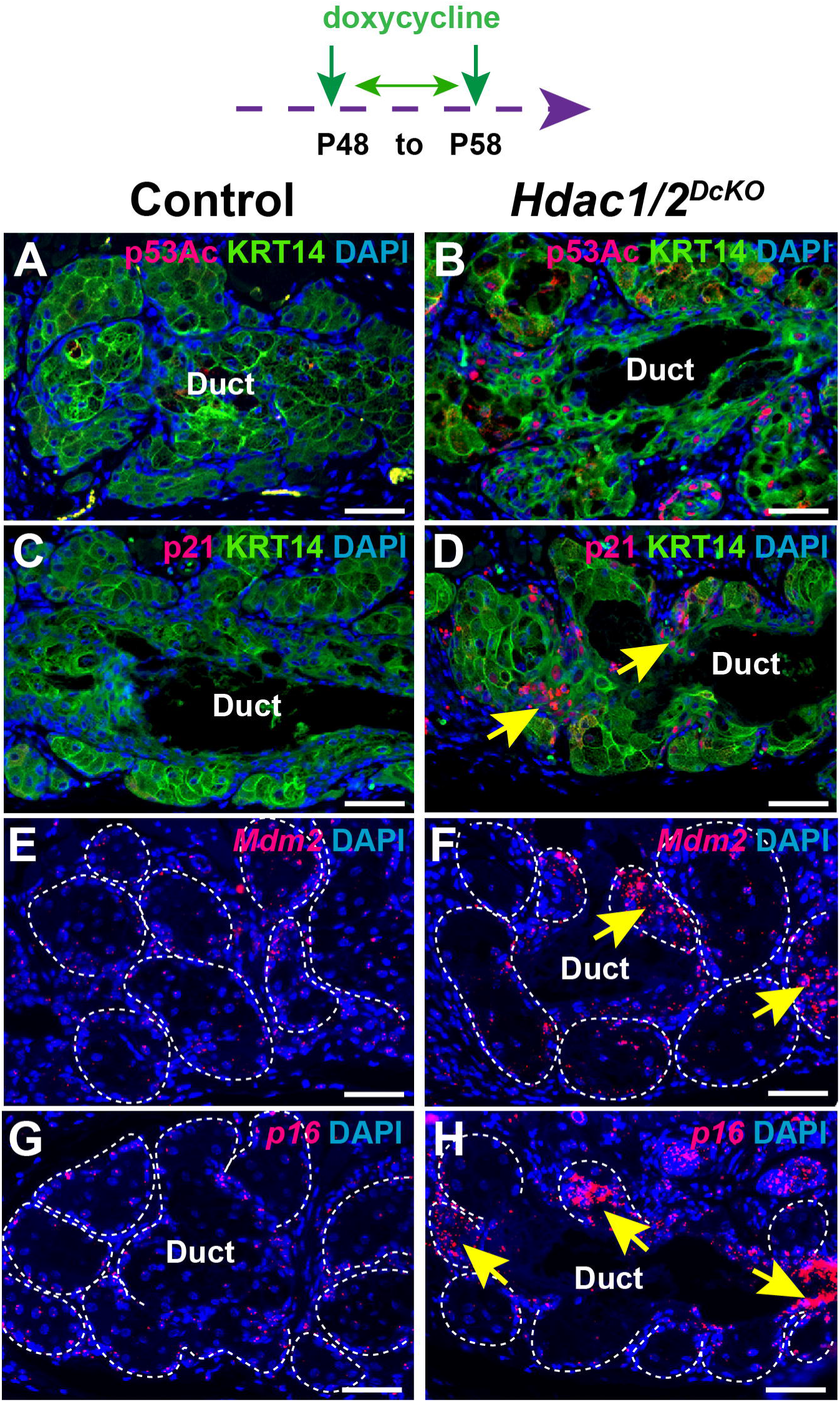
HDAC1/2 deficiency causes elevated levels of p53 acetylation and p53 target gene expression in MG. (A-D) IF reveals elevated levels of p53Ac (A, B) and p21 (C, D) in HDAC1/2-deficient MGs (B, D) compared with littermate controls (A, C). (E-H) RNAscope assays show increased expression levels of *Mdm2* (E, F) and *p16* (G, H) in HDAC1/2-deficient MGs (F, H, yellow arrows) compared with littermate controls (E, G). White dashed lines delineate MG acini. Scale bar: 50μm.

### The functions of HDAC3 in the adult MG are distinct from those of HDAC1/2

HDAC3 uniquely associates with the SMRT/N-CoR corepressor complex, suggesting that its functions in the adult MG may differ from those of HDAC1/2. To begin to explore this, we used IF to examine HDAC3 expression in adult MGs and found that it is expressed ubiquitously in MG epithelial cells (Fig. 4A). To reveal the functions of HDAC3 in MG homeostasis, we induced epithelial *Hdac3* deletion in *K5-rtTA tetO-Cre Hdac3^fl/fl^* (*Hdac3^cKO^*) mice (Fig. 4B). Histological examination after 5 weeks of induction revealed pronounced atrophy within the acini and dilation of ducts in HDAC3-deficient MGs (Fig. 4C-F). Depletion of meibocytes and ductal dilation were confirmed by IF for PLIN2 and KRT6A (Fig. 4G and 4H, white arrows). Unlike in HDAC1/2-deficient MGs, the ducts within HDAC3-deleted MGs lacked KRT10 expression (Fig. 4I and 4J), CD45+ immune cells were absent from the acini of HDAC3-deficient MGs (Fig. 4K and 4L). Thus, HDAC3-deficient MGs have phenotypes markedly different from those of HDAC1/2 deleted MGs, in line with distinct functions for HDAC3 and HDAC1/2 in adult MGs.

**Figure 4.**
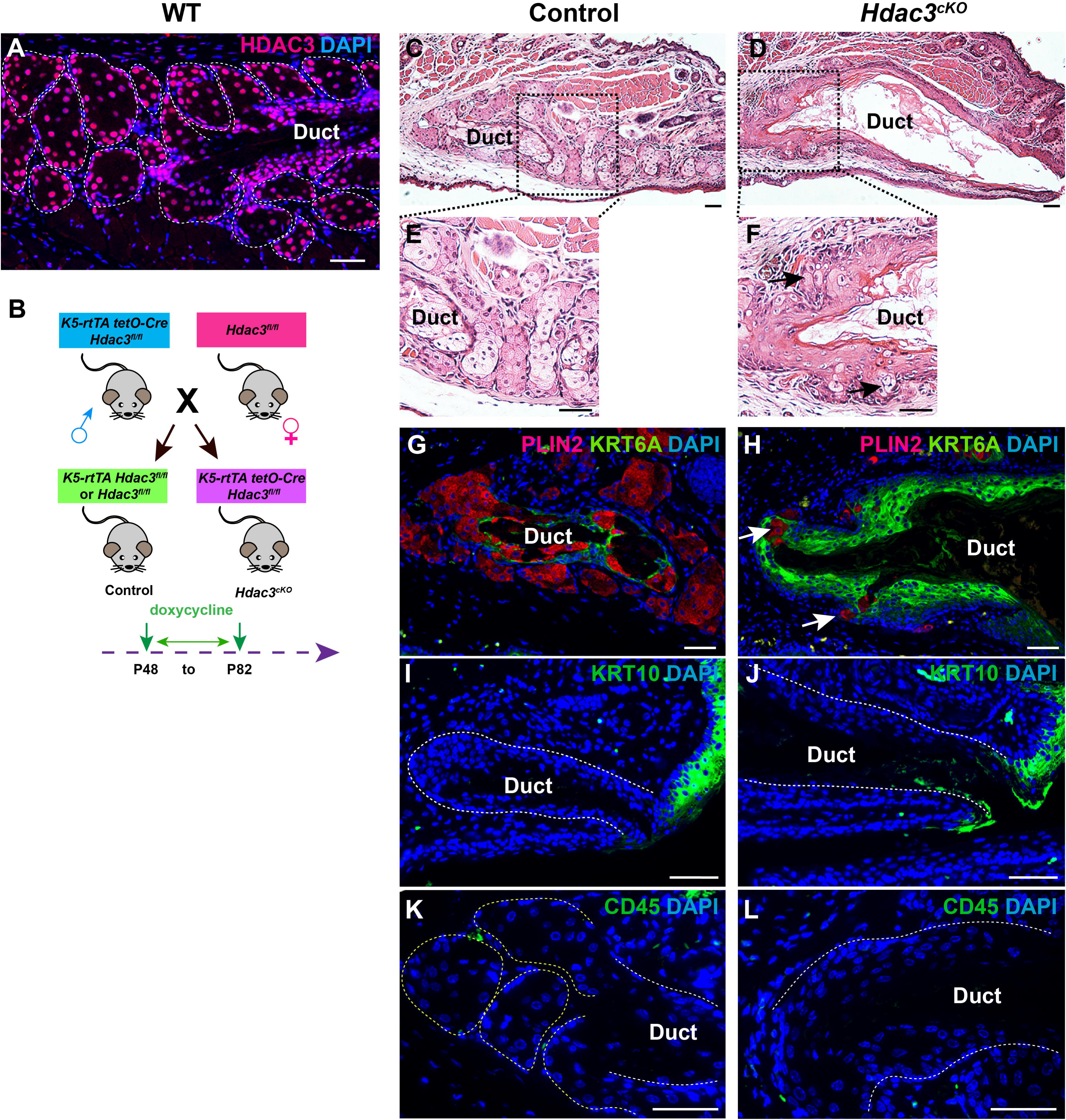
HDAC3-deficient MG exhibit atrophic acini and dilated central ducts. (A) HDAC3 is expressed ubiquitously in WT MGs. (B) Schematic depicting the mouse breeding strategy and experimental setup. (C, D) H&E staining reveals decreased acinar size and dilated ducts in HDAC3-deficient (D) compared with control (C) MG. (E, F) Enlarged views of the boxed regions in (C) and (D), respectively. Black arrows in (F) indicate miniaturized acini. (G, H) IF staining reveals expanded KRT6A expression and reduced numbers of PLIN2+ meibocytes in HDAC3-deficient MGs (H, white arrows) compared with control MG (G). (I-L) IF staining reveals absence of KRT10 expression (I, J) or CD45+ immune cell infiltration (K, L) in control (I, K) and HDAC3-deleted (J, L) MG ducts. White dashed lines delineate the MG duct and acini. Scale bars: 50μm.

### HDAC3 is required for acinar basal cell differentiation

To evaluate the proximal consequences of HDAC3 deletion on MG homeostasis, we analyzed MGs from control and *Hdac3^cKO^* mice after 10 days of doxycycline treatment. At this stage, the HDAC3-deficient MGs retained a relatively normal histological appearance, although preliminary signs of degeneration were noticeable in both the MG acinus and duct (Fig. 5A and 5B, black arrows). IF analysis confirmed efficient deletion of HDAC3 in MG epithelia (Fig. 5C and 5D). Strikingly, expression of KRT5, which is primarily limited to the basal layer of the acinus in control MGs (white arrows, Fig. 5E), was expanded towards the center of HDAC3-depleted MG acini (yellow arrows, Fig. 5F), suggesting that differentiation of acinar basal cells was perturbed upon HDAC3 ablation. Consistent with this, we identified a substantial reduction in expression of the acinar differentiation markers FASN and PPARγ in HDAC3-deficient acini (Fig. 5G-J). These data reveal an essential role for HDAC3 in promoting acinar basal cell differentiation.

**Figure 5.**
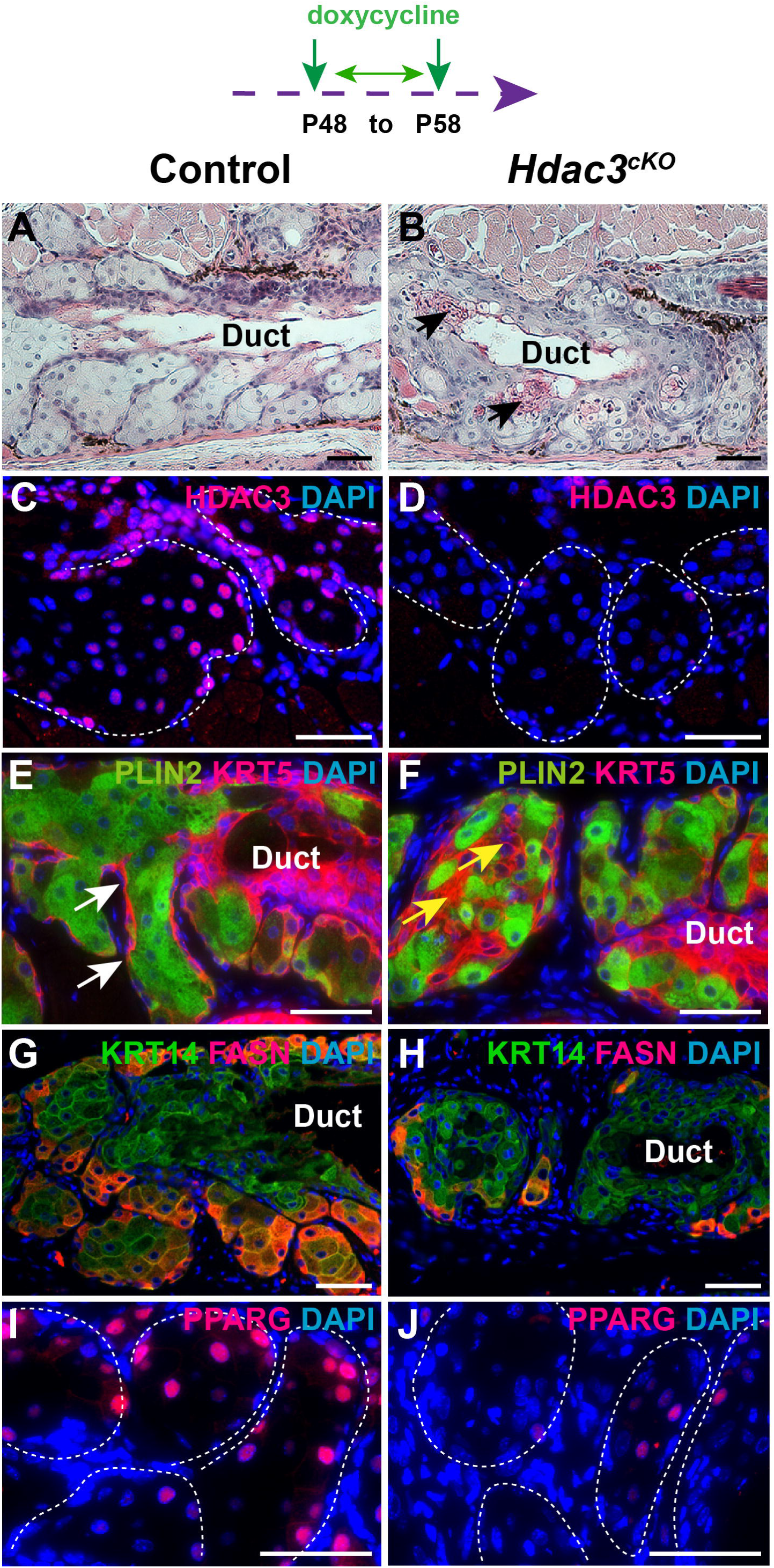
*Hdac3* deletion impairs MG acinar differentiation. (A, B) H&E staining reveals degeneration in the MG duct and acinus of *Hdac3^cKO^* mice (B, black arrows) compared with control (A) after 10 days of doxycycline treatment. (C, D) IF for HDAC3 reveals efficient deletion in *Hdac3^cKO^* MG (D) compared with control (C). (E, F) IF reveals KRT5 expression in MG duct and in the basal layer of control MG acini (E, white arrows), and expansion of KRT5+/PLIN2- cells into the center of acini in *Hdac3^cKO^* MG (F, yellow arrows). (G-I) IF data demonstrate downregulation of FASN (G, H) and PPARγ (I, J) expression in HDAC3-deleted acini (H, J) compared with littermate controls (G, I). White dashed lines delineate MG acini. Scale bars: 50μm.

### HDAC3 suppresses acinar basal cell proliferation

Expansion of HDAC3-deleted acinar basal cells suggested that HDAC3 may function to suppress basal cell proliferation. To test this, we examined expression of the proliferation marker Ki-67 and the proliferation regulator *Ccnd1*. These experiments revealed a statistically significant increase in the percentage of Ki-67+ basal cells (Fig. 6A, 6B, and 6E), and elevated expression of *Ccnd1* (Fig. 6C and 6D), in HDAC3-deficient acini. Published data from in vitro experiments indicate that the Hedgehog (Hh) signaling pathway is required for normal levels of MG epithelial cell proliferation [41]. To test whether altered Hh signaling could potentially contribute to acinar basal cell hyperproliferation in the absence of HDAC3, we examined expression of the Hh pathway target genes *Gli1* and *Ptch1*. Interestingly, expression of *Gli1* (Fig. 6F and 6G) and *Ptch1* (Fig. 6H and 6I) was increased in HDAC3-deficient MG acini compared with controls. In line with this, the Hh target gene KRT17, which is primarily expressed in MG duct and absent from the center of the acinus in control samples (Fig. 6J), was ectopically expressed in HDAC3-deficient acini (Fig. 6K, yellow arrows). These data are consistent with a model in which HDAC3 suppresses acinar basal cell proliferation in part by dampening Hh signaling pathway activity.

**Figure 6.**
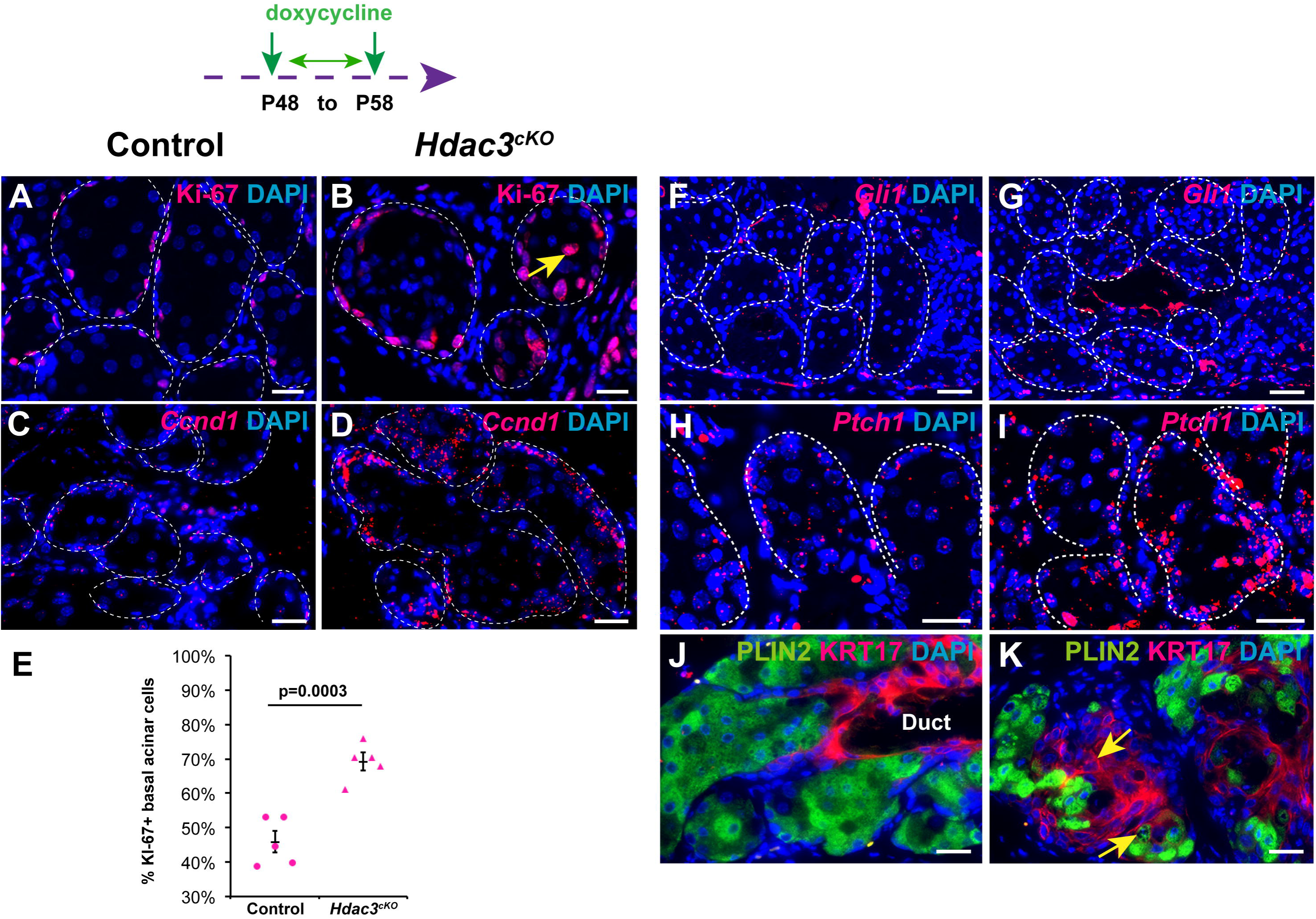
HDAC3 negatively regulates acinar basal cell proliferation. (A, B) The number of Ki-67+ cells in the basal layer of MG acini is increased 10 days after inducing HDAC3 deletion (B) compared with littermate control (A). Notably, Ki-67+ cells are observed in the center of HDAC3-deleted acini (B, yellow arrow) as well as in the basal layer. (C, D) RNAscope data reveal increased expression of *Ccnd1* in the acini of HDAC3-deficient (D) compared with control (C) MG. (E) Quantification of proliferation in basal acinar cells of control and HDAC3-deleted MGs. Five pairs of control and *Hdac3^cKO^* mice were analyzed. Error bars indicate SEM. (F-I) RNAscope data reveal increased expression of *Gli1* (F, G) and *Ptch1* (H, I) in the acini of HDAC3-deficient MGs (G, I) compared with controls (F, H). (J, K) IF for KRT17 and PLIN2 reveals their expression confined to duct and acini, respectively, in control MG (J); in *Hdac3^cKO^*MG, PLIN2 expression is reduced, and KRT17 is ectopically expressed in MG acini (H, yellow arrows). White dashed lines delineate MG acini. Scale bars: 25μm.

### HDAC3 prevents apoptosis and DNA damage and suppresses p53 pathway signaling in adult MG

To determine whether HDAC3 deletion impacts MG cell survival, we performed TUNEL assays. The results showed that HDAC3 ablation triggered ectopic apoptosis in both the MG duct (yellow arrows) and acini (white arrows) (Fig. 7A and 7B). IF analysis for pH2A.X revealed that DNA damage was limited to fully differentiated meibocytes in control MG acini (Fig. 7C, yellow arrows), but in HDAC3-deficient MG was also observed in acinar basal cells (pink arrows in Fig. 7D) and duct basal cells (white arrows in Fig. 7D). Levels of acetylated p53 (p53Ac) (Fig. 7E and 7F) and the p53 target genes p21 (Fig. 7G and 7H) and *Mdm2* (Fig. 7I and 7J) were elevated in HDAC3-deficient compared with control MGs. In contrast to HDAC1/2-deleted MGs, HDAC3 deletion exhibited minimal impact on *p16* expression, as revealed by the results of RNAscope analysis (Fig. 7K and 7L). Collectively, these findings suggest that HDAC3 promotes cell survival within MGs in part by causing p53 deacetylation while leaving *p16* expression largely unaffected.

**Figure 7.**
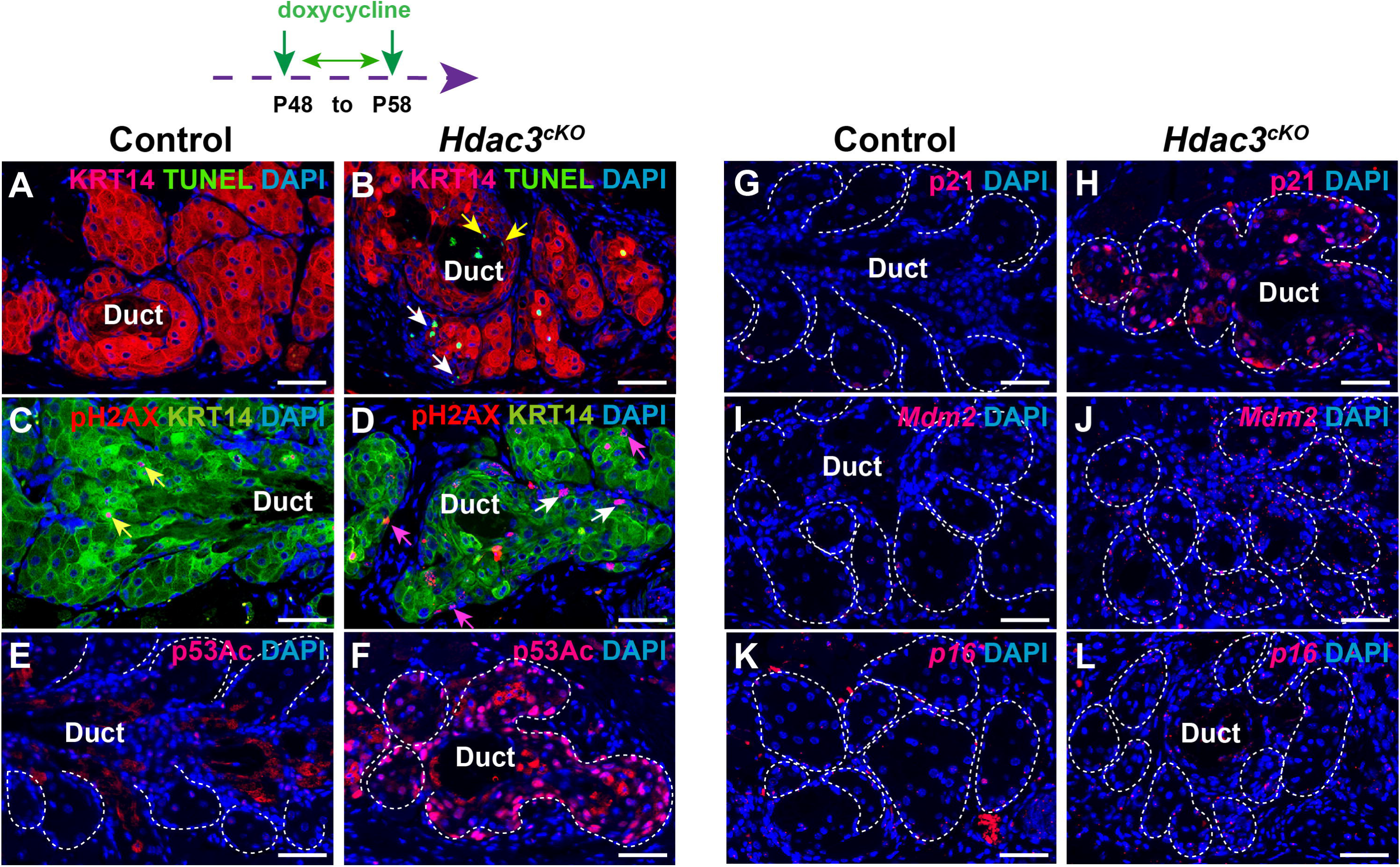
*Hdac3* deletion causes ectopic apoptosis, increased DNA damage, and activation of the p53 pathway in MGs. (A, B) TUNEL assays reveal ectopic apoptosis in the acini (B, white arrows) and duct (B, yellow arrows) of HDAC3-deficient (B) compared with control (A) MG 10 days after initiating doxycycline treatment. (C, D) IF data for pH2A.X indicate that DNA damage is confined to mature meibocytes in control MG (C, yellow arrows). HDAC3- deficient MGs display increased DNA damage in the basal layer of the acinus (D, pink arrows) and duct (D, white arrows). (E-H) IF data indicate low levels of p53Ac (E, F) and p21 (G, H) expression in control MGs (E, G) and elevated levels of p53Ac and p21 in HDAC3-deficient MGs (F, H). (I-L) RNAscope data indicate low expression of *Mdm2* (I) and *p16* (K) in control MGs. In HDAC3-deficient MGs, *Mdm2* expression is increased (J) but *p16* expression is not significantly altered (L) compared with controls. White dashed lines delineate the MG acini. Scale bars: 50μm.

## Discussion

The mechanisms governing MG homeostasis are poorly understood, particularly at the epigenetic level. In this study, we provide definitive genetic evidence that epigenetic control orchestrated by HDAC1/2 and HDAC3 is vital for maintaining the homeostasis of adult MG.

Although HDAC1 and HDAC2 possess distinct functions, they exhibit redundancy in fulfilling their roles in several different cellular contexts [24, 37, 42–46]. In line with this, we did not observe any significant morphological alterations in the MGs of *Hdac1^cKO^ Hdac2^cHet^* or *Hdac1^cHet^ Hdac2^cKO^* mice following induction of gene deletion in adult mice, whereas the MGs of *Hdac1^cKO^ Hdac2^cKO^*mice exhibited strong phenotypes including decreased epithelial cell proliferation and survival in the same time frame. These data suggest that *Hdac1* and *Hdac2* function redundantly in adult MG homeostasis.

These findings contrast with the observation that constitutive homozygous deletion of *Hdac1* combined with heterozygous deletion of *Hdac2* in *K14-Cre Hdac1^fl/fl^ Hdac2^fl/+^* mice during MG development results in the formation of hyperplastic MGs [36]. Taken together, these data suggest that *Hdac1* plays some essential roles in MG development that cannot be substituted by *Hdac2*, whereas these two genes are fully redundant during adult MG homeostasis. However, it remains possible that the different phenotypic outcomes of constitutive versus inducible deletion are due to use of the *K14-Cre* versus *K5-rtTA tetO-Cre* drivers, to differences in mouse strain background between the published study and that reported here, or to the relatively limited timeframe between gene deletion and tissue analysis used in the current study.

HDAC1/2 are known to promote proliferation and suppress apoptosis in a range of different tissues [16]. In adult epidermis, HDAC1/2 regulate the proliferation and survival of keratinocytes via p53 and p16 [25]. In line with this, our investigations revealed increased acetylated p53 levels and p16 expression in HDAC1/2-depleted MG cells. These results suggest the existence of overlapping mechanisms of action of HDAC1/2 in keratinocytes and MG epithelial cells. Given the diverse spectrum of targets modulated by HDAC1/2, it will be interesting in the future to identify additional downstream factors governed by HDAC1/2 in MG homeostasis and to determine the extent to which these also operate in epidermal keratinocytes.

In contrast to the proliferation-enhancing role of HDAC1/2, HDAC3 appears to play an opposing function by negatively modulating the proliferation of acinar basal cells within MGs. This role diverges from HDAC3’s function in some cellular contexts where it stimulates proliferation [47–50], highlighting a distinctive role of HDAC3 in governing MG acinar basal cell division and limiting the expansion of KRT5+ basal cells. Interestingly, we found that HDAC3-deficient acinar basal cells exhibit elevated expression levels of *Gli1*, *Ptch1* and KRT17, which are targets of the Hh signaling pathway. In vitro experiments have revealed that Hh signaling is required for MG cell proliferation [41]. Taken together, these observations suggest that HDAC3 may repress proliferation within the basal layer of the acini via modulation of the Hh pathway.

In addition to causing expansion of MG basal cells, HDAC3 deficiency perturbs meibocyte differentiation. Specifically, expression of PPARγ, a key regulator promoting meibocyte differentiation [51, 52], was markedly downregulated in HDAC3-depleted MG acini. In line with this, HDAC3 positively regulates *Pparg* expression in lung alveolar macrophages by binding to *Pparg* enhancers [53]. By contrast, in adipocytes HDAC3 deacetylates PPARγ resulting in its downregulation [54]. These data indicate that the influence of HDAC3 on PPARγ expression and activity is contingent on the specific cellular context.

Despite marked differences in the MG phenotypes caused by inducible deletion of *Hdac3* versus *Hdac1/2*, in both cases we observed activation of the p53 pathway through elevated p53 acetylation, consistent with previous observations in several different cell types [55, 56]. Our data indicate that p53 serves as a shared target in MG epithelial cells, regulated by both HDAC1/2 and HDAC3. However, the comparatively milder apoptosis and MG degeneration observed in HDAC3-compared with HDAC1/2-deficient mice suggest that HDAC1/2 may employ additional mechanisms to bolster the survival of MG cells. In line with this, the expression of *p16* is upregulated in HDAC1/2-depleted MGs but appears unaffected by HDAC3 deletion.

In summary, our findings reveal that HDAC1/2 and HDAC3 play distinct, and in some cases opposing, roles in maintaining adult MG homeostasis. Thus, the effects of using broadly acting HDAC inhibitors on both unaffected MG tissue and MG tissue affected by diseases such as DED are likely to be complex. Our results suggest a need to develop inhibitors that can modulate the effects of individual HDAC family members, and the importance of identifying additional, disease-specific, therapeutic targets.

## Acknowledgments

We thank Adam Glick for *K5-rtTA* mice; Eric N. Olson for *Hdac1^fl/fl^* and *Hdac2^fl/fl^* mice; and Mitchell A Lazar for *Hdac3^fl/fl^* mice. This work was supported by NIAMS/NIH grants RO1 AR081322 and R37 AR047709 (SEM) and a pilot grant from the NIAMS/NIH-supported Skin Biology and Diseases Resource-based Center (SBDRC) P30 AR079200 (XZ).

## Author contributions

Conceptualization: XZ, MX, SEM; Formal Analysis: XZ; Funding Acquisition: XZ, SEM; Investigation: XZ; Methodology: XZ; Project Administration: SEM; Resources: XZ, SEM; Supervision: SEM; Validation: XZ; Visualization: XZ; Writing - Original Draft Preparation: XZ; Writing - Review and Editing: XZ, MX, SEM.

## Data availability statement

The authors declare that the main data supporting the findings of this study are available within the article and its Supplemental Information files. All correspondence and material requests should be addressed to Sarah E. Millar. This study includes no data deposited in external repositories.

---

**Figure S1.** Compound homozygous/heterozygous deletion of HDAC1 and HDAC2 in MG epithelium has minimal effect on MG homeostasis. (A) Schematic depicting mouse breeding strategy and experimental setup. (B-D) MG histology in control littermate (B), *Hdac1^cKO^ Hdac2^cHet^* mice (C), and *Hdac1^cHet^ Hdac2^cKO^*mice (D). (E-G) IF for HDAC1 confirming efficient deletion of HDAC1 in the MG of *Hdac1^cKO^ Hdac2^cHet^* mice (F) and maintained expression of HDAC1 in *Hdac1^cHet^ Hdac2^cKO^* mice (G), compared with littermate control (E). (H-J) IF for HDAC2 showing its expression in control (H) and *Hdac1^cKO^ Hdac2^cHet^* (I) MGs, and its efficient deletion in *Hdac1^cHet^ Hdac2^cKO^* MG (J). (K-M) Unaffected expression of PLIN2 and KRT6A in the MG of *Hdac1^cKO^ Hdac2^cHet^* mice (L) and *Hdac1^cHet^ Hdac2^cKO^* mice (M), compared with control (K). Scale bars: 50μm.

