## Supplementary figures and images for "HDAC1/2 and HDAC3 play distinct roles in controlling adult Meibomian gland homeostasis"

### Fig.S1

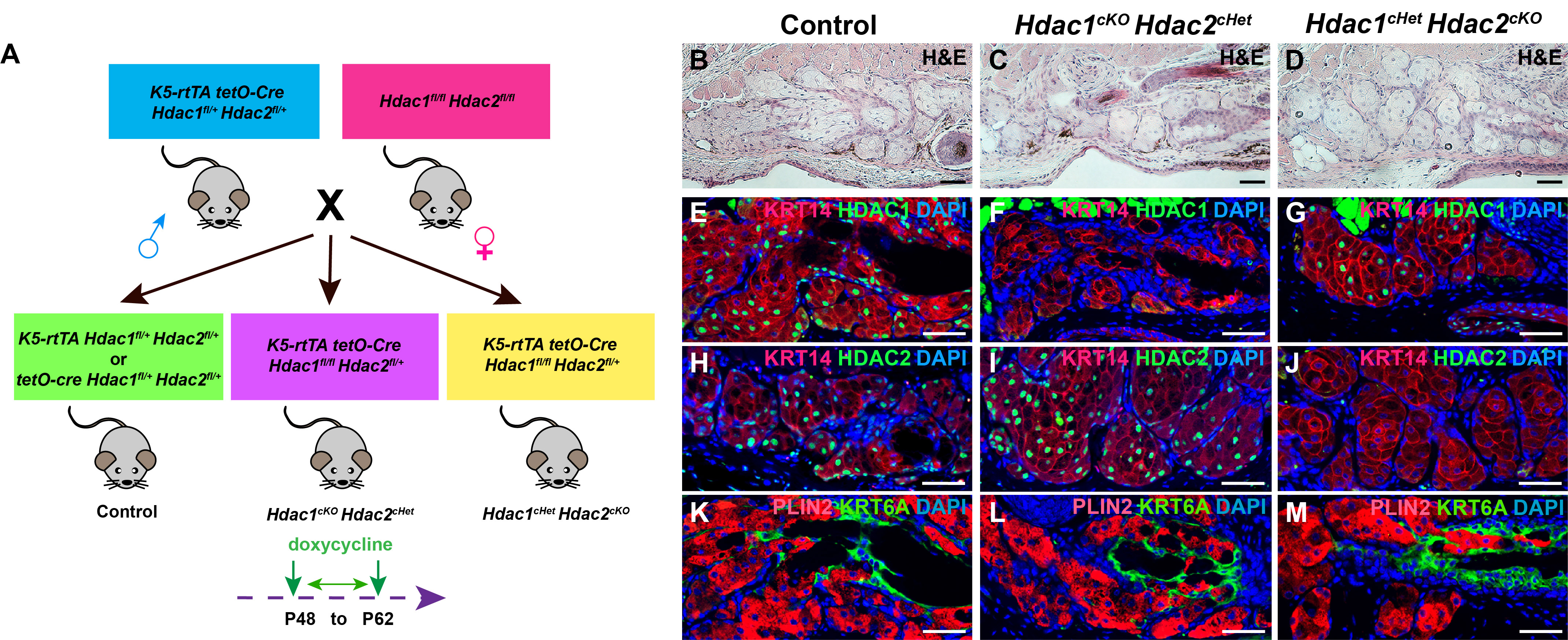
